# Detoxification of host-relevant H_2_O_2_ shapes *Pseudomonas aeruginosa* population gradients in flow

**DOI:** 10.64898/2026.05.22.727215

**Authors:** Anuradha Sharma, Alexander M. Shuppara, Joseph E. Sanfilippo

## Abstract

During infection, bacterial populations need to overcome low levels of host-generated H_2_O_2_. However, experiments almost always use high H_2_O_2_ levels that are rarely found in hosts. Here, we use microfluidics to investigate how host-relevant H_2_O_2_ impacts populations of the human pathogen *Pseudomonas aeruginosa* in flow. Using long-channel microfluidic devices, we establish that cells at the front of a population remove H_2_O_2_ and protect their downstream neighbors. Population-level protection is mediated by three OxyR-regulated scavenging systems (KatA, KatB, AhpCF), which are each sufficient to detoxify host-relevant H_2_O_2_. Mutants lacking all three systems are sensitive to H_2_O_2_ but can be cross-protected by resistant cells when co-cultured. Cross-protection is abolished in higher flow regimes, where H_2_O_2_ is delivered faster than cells can remove it. Our results demonstrate how local detoxification provides global protection, which results in spatial gradients across bacterial populations in flow. Together, our findings highlight how biological, chemical, and physical factors collectively determine the fate of bacterial populations in host-relevant environments.

## Introduction

A bacterium’s life is shaped by its physical and chemical environment. During colonization of the bloodstream, lungs, or urinary tract, bacteria are exposed to host-generated fluid flow (1–3). Using microfluidic approaches, studies have shown that flow can impact bacterial behavior in several ways (4). While the force associated with flow can impact motility and adhesion (5–9), flow can also impact bacteria in a force-independent manner (10–14). By facilitating chemical transport, flow can replenish or wash away small molecules that impact growth and cell-cell communication. For example, flow promotes growth in nutrient-limited environments (14), enhances antimicrobial effectiveness (10, 11, 13), and inhibits quorum sensing (15). As flow is a key determinant of bacterial success or failure, there is a critical need to study bacterial behavior in host-relevant flow.

During infection, bacteria are exposed to combinations of physical and chemical stressors. However, most studies examine bacterial responses to isolated stressors. During colonization of the bloodstream, lungs, or urinary tract, bacteria are exposed to host-generated H_2_O_2_. Although H_2_O_2_ stress responses have been extensively studied (16, 17), nearly all experiments have used H_2_O_2_ levels much higher than those found in hosts (18–21). This has led to the conclusion that high levels (millimolar) of H_2_O_2_ are required to inhibit bacterial growth (22–26). In contrast, microfluidic experiments that combine flow and H_2_O_2_ revealed that host-relevant levels (micromolar) of H_2_O_2_ can inhibit bacterial growth in flow (11). Flow enhances H_2_O_2_ effectiveness by delivering H_2_O_2_ faster than cells can detoxify it (13), highlighting how physical and chemical stressors can have synergistic effects.

When bacterial cells grow into a population, chemical gradients naturally emerge as signaling molecules, metabolites, and nutrients are secreted or consumed. Flow-mediated transport of small molecules shapes local chemical gradients across populations. For example, flow can inhibit quorum sensing of upstream cells by washing away autoinducers that ultimately activate quorum sensing of downstream cells (15). Similarly, flow can transport metabolic by- products from upstream to downstream species, creating spatial structure within a mixed population (27). Alternatively, flow can replenish scarce nutrients as they are consumed, allowing upstream cells to grow better than downstream cells (14). As flow can replenish H_2_O_2_ faster than bacteria can detoxify it, there is a prime opportunity to study the spatial structure of bacterial populations treated with H_2_O_2_ in flow.

Here, we use microfluidics to examine how flow and H_2_O_2_ impact populations of the human pathogen *Pseudomonas aeruginosa*. Using host-relevant flow, we learn that *P. aeruginosa* populations can detoxify host-relevant H_2_O_2_ via three OxyR-regulated scavenging systems (KatA, KatB, AhpCF). Detoxification of H_2_O_2_ results in spatial gradients, where cells at the start of a microfluidic channel promote growth of their downstream neighbors. Surprisingly, we discover that M9 minimal medium contains low levels of H_2_O_2_ that are sufficient to block growth of an H_2_O_2_-sensitive mutant. By mixing sensitive and resistant strains together, we show that bacterial cells in a population can protect one another from H_2_O_2_. Collectively, our results demonstrate how *P. aeruginosa* cells defend against host-relevant H_2_O_2_ and reveal that survival strategies benefiting individual cells can provide communal protection as a byproduct.

## Results

As a foundation for our study of bacterial H_2_O_2_ defenses, we first compared the H_2_O_2_ concentrations found in the human body to those typically used in experiments. While host environments such as the bloodstream, urinary tract, and lungs typically have 1-25 µM H_2_O_2_ (28–30), experiments have typically used 1-10 mM (19–21), which is approximately 1,000 times higher (Figure 1A). Motivated by this discrepancy, we used host-relevant H_2_O_2_ concentrations to characterize the key molecular players that underlie the *P. aeruginosa* H_2_O_2_ response. The central protein is the H_2_O_2_ sensor OxyR, which directly senses H_2_O_2_ by forming a disulfide bond between two conserved cysteine residues (31, 32). Once activated, OxyR transcriptionally activates genes that encode three scavenging systems: a primary catalase (KatA), a secondary catalase (KatB), and two proteins that act as alkyl hydroperoxide reductases (AhpC and AhpF) (22, 33) (Figure 1B).

**Figure 1:**
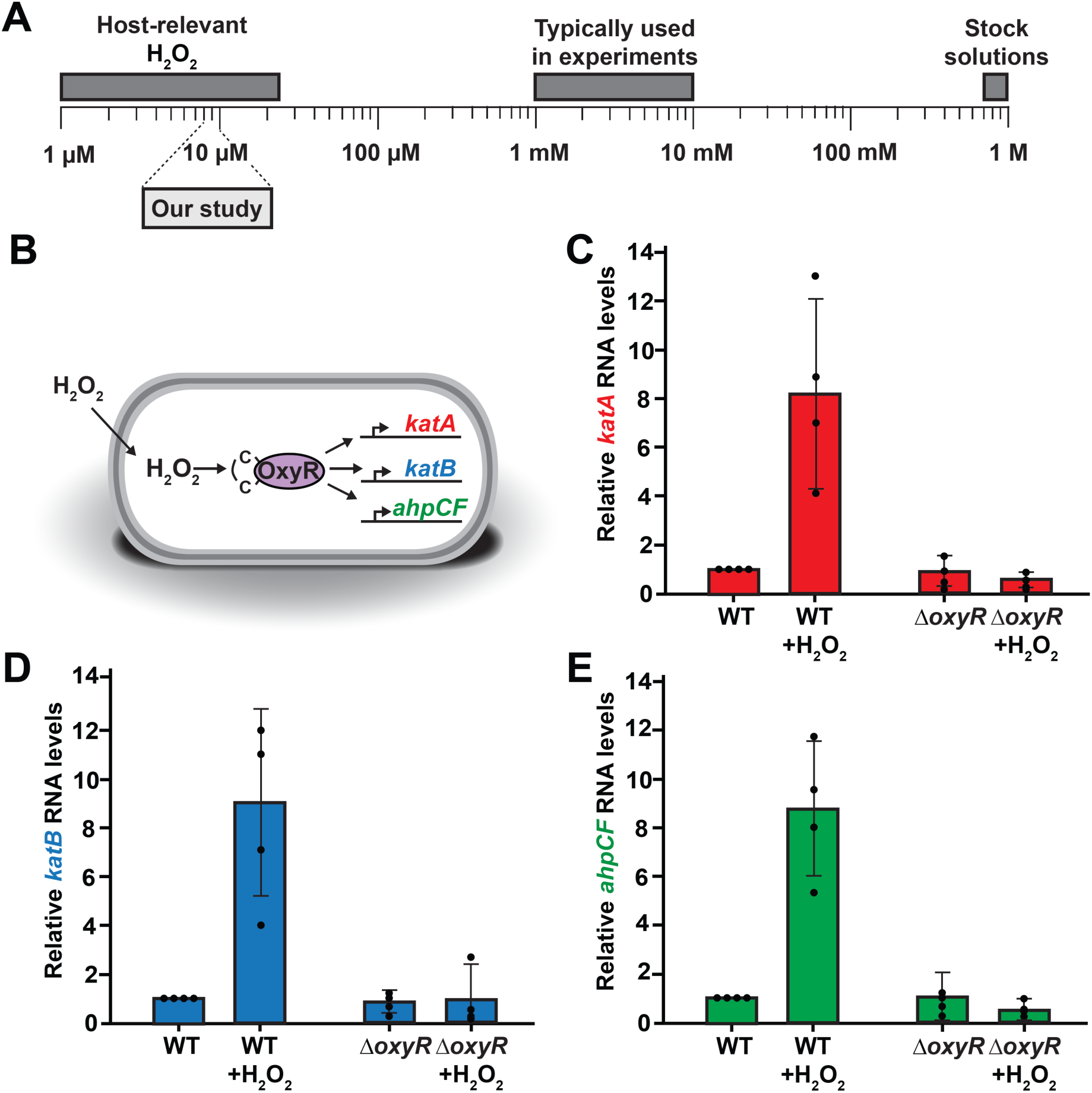
Host-relevant H_2_O_2_ induces three OxyR-regulated scavenging systems. **(A)** Representation of H_2_O_2_ concentrations. The right side of the scale represents concentrations found in commercially available stock solutions (∼1 M) and those typically used in laboratory experiments (1-10 mM). The left side of the scale represents the host-relevant H_2_O_2_ levels in human tissues (1-25 µM). Experiments in our study were conducted at host-relevant H_2_O_2_ levels (8-10 µM). **(B**) Model of OxyR-mediated gene expression in response to H_2_O_2_ stress. OxyR directly senses H_2_O_2_ through reactive cysteine residues and activates transcription of H_2_O_2_ sensitive genes (*katA*, *katB*, *ahpCF*). **(C**) *katA* RNA levels in WT and Δ*oxyR* cells from a qRT-PCR experiment with and without 8 μM H_2_O_2_. **(D)** *katB* RNA levels in WT and Δ*oxyR* cells from a qRT-PCR experiment with and without 8 μM H_2_O_2_. **(E)** *ahpCF* RNA levels in WT and Δ*oxyR* cells from a qRT-PCR experiment with and without 8 μM H_2_O_2_. Error bars represent SD of three biological replicates. P-value for gene induction in response to 8 µM H_2_O_2_ is < 0.05 (calculated using t-test).

To test the impact of host-relevant H_2_O_2_ on OxyR-dependent regulation, we treated wild-type cells with 8 μM H_2_O_2_ and measured transcript levels with qRT-PCR. For wild-type cells, expression of *katA*, *katB*, and *ahpCF* increased approximately 8-10-fold after 8 μM H_2_O_2_ treatment, demonstrating that host-relevant H_2_O_2_ levels are sufficient to induce the H_2_O_2_ response (Figure 1). In contrast, *ΔoxyR* cells exhibited no induction of *katA*, *katB*, and *ahpCF* expression, indicating that OxyR controls the H_2_O_2_ response to low H_2_O_2_ doses. To capture the relative expression levels of these genes, we reanalyzed previous RNA-sequencing data and ranked all the genes in the genome by transcript levels. While *katA*, *katB*, and *ahpCF* were all highly induced following 8 µM H_2_O_2_ treatment, the gene encoding the primary catalase KatA was already very highly expressed in the absence of added H_2_O_2_ (Figure S1). Together, our results indicate that OxyR induces expression of all three scavenging systems in response to host-relevant H_2_O_2_.

How do H_2_O_2_ scavenging systems impact population growth in flow? To measure the growth across *P. aeruginosa* populations, we custom-fabricated 27-cm-long microfluidic devices that allowed us to capture population growth with spatial resolution. We seeded devices with cells and used a syringe pump to introduce M9 minimal medium supplemented with glucose and 8 µM H_2_O_2_ at host-relevant shear rate of 240 s^-1^. Shear rate is a force-independent measure of fluid flow that is calculated using the flow rate and channel dimensions (4). By tracking and quantifying growth of single cells, we observed normal growth for wild-type cells throughout the channel (Figure 2A and 2B). In contrast, a triple mutant lacking *katA, katB,* and *ahpCF* exhibited no growth throughout the channel (Figure 2A and 2B). These results demonstrate that H_2_O_2_ scavenging systems are required for *P. aeruginosa* population growth in flowing environments with host-relevant H_2_O_2_ stress.

**Figure 2:**
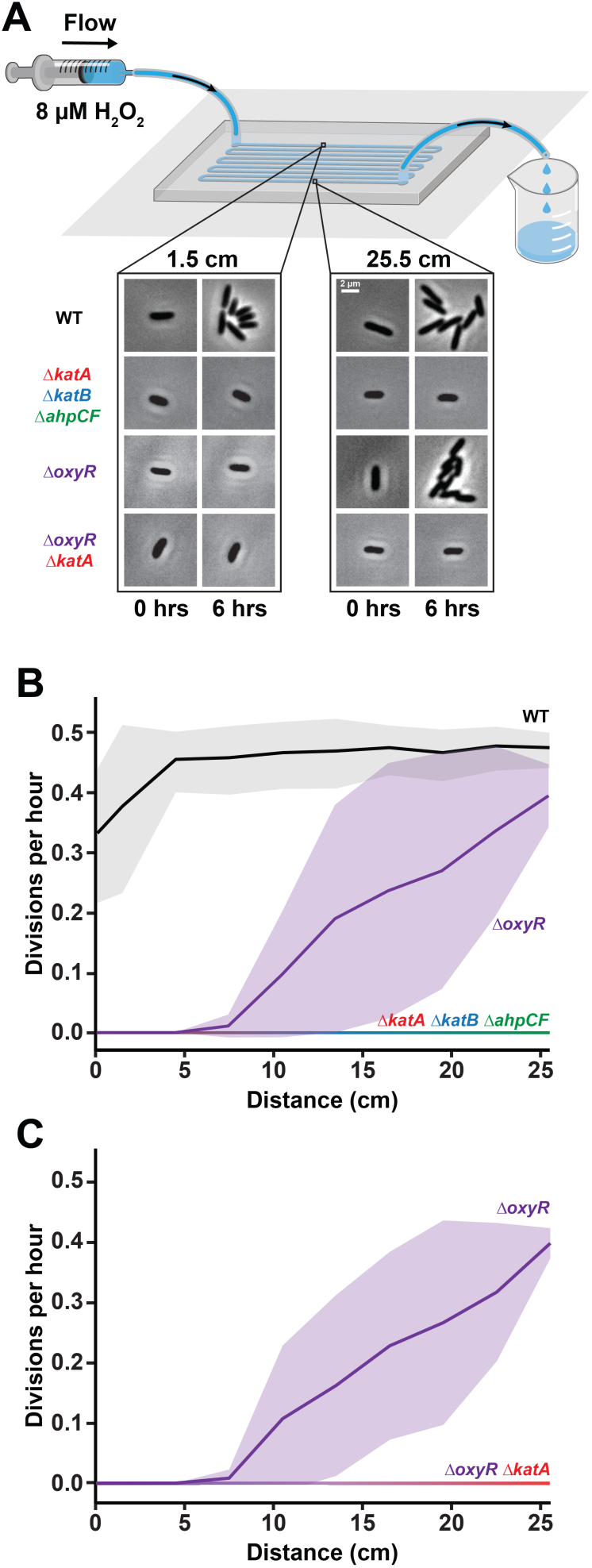
H_2_O_2_ scavenging systems shape spatial growth gradients of *P. aeruginosa*. **(A)** Experimental setup with a long-channel microfluidic device. Flow was generated with a syringe pump, and images were taken at various distances across the channel. Cell images representative of growth across a 27 cm microfluidic channel over 6 hours at 8 µM H_2_O_2_ at a shear rate of 240 s^-1^. WT cells divide every two hours. *ΔoxyR* cells do not grow in the beginning of the channel (0.1 cm) and grow by the end of the channel (25.5 cm). *ΔkatA ΔkatB ΔahpCF* and *ΔoxyR ΔkatA* cells do not grow throughout the channel. **(B**) Quantification of growth across long channel for WT and *ΔkatA ΔkatB ΔahpCF* cells. **(C)** Quantification of growth across long channel for *ΔoxyR* and *ΔoxyR ΔkatA* cells. Quantification shows the average and shaded regions show SD of three biological replicates.

To examine how OxyR impacts bacterial populations, we measured the growth of *ΔoxyR* cells in long microfluidic devices. Surprisingly, *ΔoxyR* cells treated with 8 µM H_2_O_2_ at a shear rate of 240 s^-1^ exhibited a gradient of growth across the channel. Cells failed to grow in the upstream portion of the channel but exhibited robust growth similar to wild-type in the downstream portion of the channel (Figure 2A, 2C). The fact that *ΔoxyR* cells generated spatial gradients but *ΔkatA ΔkatB ΔahpCF* cells did not led us to question what is different between these strains. As *katA* is highly expressed even in the absence of H_2_O_2_ (Figure S1), we hypothesized that uninduced *katA* was responsible for *ΔoxyR* gradients. Consistent with our hypothesis, a *ΔoxyR ΔkatA* mutant did not grow throughout the channel and did not form a spatial gradient. Together, our results indicate that OxyR-dependent regulation of H_2_O_2_ scavenging systems shapes the spatial growth patterns of *P. aeruginosa* cells in flow.

While performing experiments in long microfluidic channels, one of our negative controls generated an unexpected result. We observed that *ΔoxyR* cells exposed to M9 minimal medium without added H_2_O_2_ did not grow at the beginning of a microfluidic channel (Figure 3A). We did not expect this result because *ΔoxyR* cells should grow normally in the absence of H_2_O_2_. As wild-type cells grew normally in the same conditions (Figure 3A), we hypothesized that M9 minimal medium has trace amounts of H_2_O_2,_ and we treated our medium with catalase to remove all potential H_2_O_2_. Supporting our hypothesis, *ΔoxyR* cells exhibited normal growth in catalase-treated M9 minimal medium (Figure 3B). Consistent with our previous results, *ΔoxyR* cells generated spatial gradients in M9 minimal medium without added H_2_O_2_ (Figure 3A), and these gradients were not observed with our catalase-treated medium (Figure 3B). To further establish that M9 minimal medium contains H_2_O_2,_ we quantified the H_2_O_2_ levels using a fluorometric assay. We observed that M9 minimal medium has approximately 0.5 µM H_2_O_2_ (Figure 3C). This unexpected observation supported our results that very low micromolar levels of H_2_O_2_ doses are sufficient to shape population growth gradients in flow.

**Figure 3:**
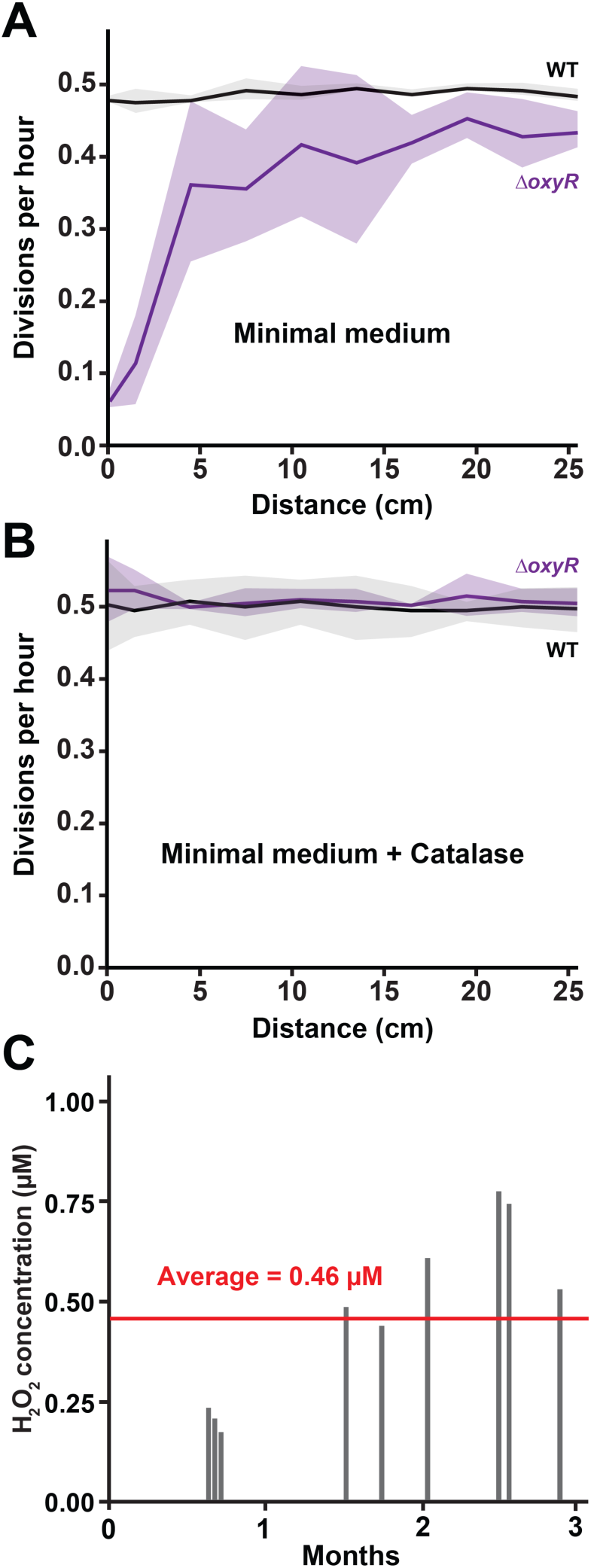
H_2_O_2_ from minimal media creates a gradient of *P. aeruginosa* growth. **(A)** Quantification of growth in M9 minimal medium with no added H_2_O_2_ at a shear rate of 240 s^-1^. WT cells exhibit normal growth while *ΔoxyR* cells exhibit a gradient of growth across the channel. **(B**) Quantification of growth in M9 minimal medium supplemented with catalase at a shear rate of 240 s^-1^. WT cells and *ΔoxyR* cells exhibit normal growth throughout the channel. **(C)** H_2_O_2_ measurements in M9 minimal medium over months. Shaded regions show SD of three biological replicates.

As H_2_O_2_ detoxification shapes spatial growth patterns, we aimed to further understand the contributions of the three H_2_O_2_ scavenging systems. KatA and KatB each function as catalases to break down H_2_O_2_ into water and O_2_, while AhpC and AhpF work together to reduce H_2_O_2_ to water using NADH as a reductant (Figure 4A). To dissect the role of each scavenging system, we first measured the ability of single mutant strains to scavenge H_2_O_2_ from a test tube. As *ΔkatA*, *ΔkatB*, and *ΔahpCF* single mutants all scavenged H_2_O_2_, we concluded that they were functionally redundant in our assay (Figure 4B). Similarly, *ΔkatA ΔkatB*, *ΔkatA ΔahpCF*, and *ΔkatB ΔahpCF* double mutants all scavenged H_2_O_2_ (Figure S3), reinforcing the conclusion that the three scavenging systems are functionally redundant in our assay.

**Figure 4:**
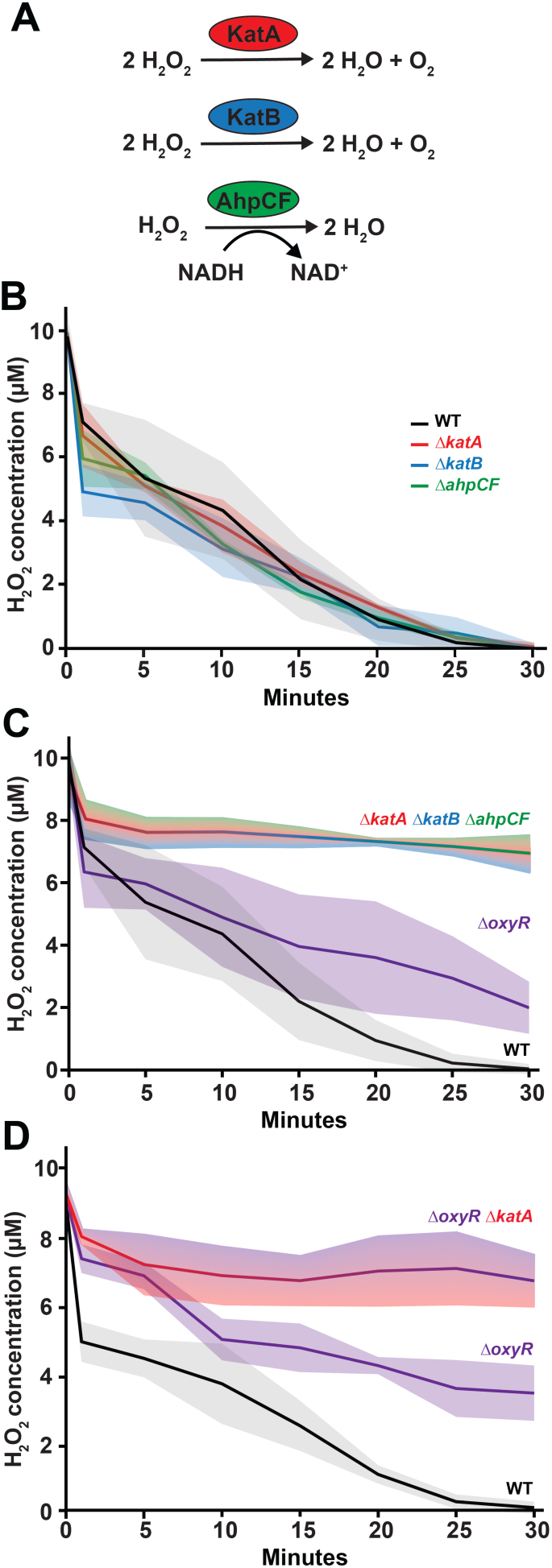
*P. aeruginosa* H_2_O_2_ scavenging systems remove H_2_O_2_ over time. **(A)** Biochemical reactions carried out by H_2_O_2_ scavenging enzymes. KatA, KatB, and AhpCF break down H_2_O_2_. **(B)** H_2_O_2_ scavenging by WT, *ΔkatA*, Δ*katB*, Δ*ahpCF* cells. All strains scavenge 10 µM H_2_O_2_ in 30 minutes. **(C**) H_2_O_2_ scavenging by WT and *ΔkatA ΔkatB ΔahpCF* cells. Δ*katA ΔkatB ΔahpCF* cells are impaired at scavenging. **(D)** H_2_O_2_ scavenging by WT, *ΔoxyR* and *ΔoxyR ΔkatA* cells. *ΔoxyR* cells are slower in scavenging that WT cells. Δ*oxyR ΔkatA* cells are impaired at scavenging. All scavenging assays are performed at a 1:10 dilution of 0.2 OD cells for 30 minutes. Quantification shows the average and shaded regions show SD of three biological replicates.

To further examine the role of the three scavenging systems, we compared H_2_O_2_ scavenging by wild-type and *ΔkatA ΔkatB ΔahpCF* triple mutant cells. While wild-type cells scavenged H_2_O_2_, cells lacking all three systems were severely impaired at scavenging (Figure 4C). As *ΔoxyR* cells and the triple mutant cells display different growth patterns in flow, we questioned whether they differ in their ability to scavenge H_2_O_2_. *ΔoxyR* cells scavenged H_2_O_2_ more effectively than the triple mutant cells but at a slower rate than wild-type cells (Figure 4D). Notably, *ΔoxyR ΔkatA* cells were severely impaired at scavenging H_2_O_2_ (Figure 4D), confirming that uninduced *katA* was responsible for scavenging in *ΔoxyR* cells. Together, the differences in scavenging capacity coupled with spatial gradients observed in flow demonstrate that bacterial population growth is dependent on H_2_O_2_ detoxification across the microfluidic channel.

As spatial gradients depend on H_2_O_2_ detoxification, we wondered if resistant cells could protect sensitive cells in co-culture. To test our hypothesis, we subjected a co-culture of wild-type cells (90%) with triple mutant cells (10%) to 8 µM H_2_O_2_ at a shear rate of 240 s^-1^. In mono-culture experiments, wild-type cells exhibited normal growth whereas triple mutant cells did not grow throughout the channel (Figure 5A, S3). However, when wild-type cells and triple mutant cells were co-cultured, both exhibited normal growth by the end of the channel (Figure 5A, 5B). Our results demonstrated that wild-type cells scavenge H_2_O_2_ enabling growth of triple mutant cells. Similarly, co-culture experiments with wild-type and *ΔoxyR ΔkatA* cells showed that wild-type cells also supported the growth of *ΔoxyR ΔkatA* cells under H_2_O_2_ stress (Figure S4). Together, our findings support the hypothesis that resistant cells (wild-type) can protect sensitive cells (*ΔkatA ΔkatB ΔahpCF* or *ΔoxyR ΔkatA*) in co-culture.

**Figure 5:**
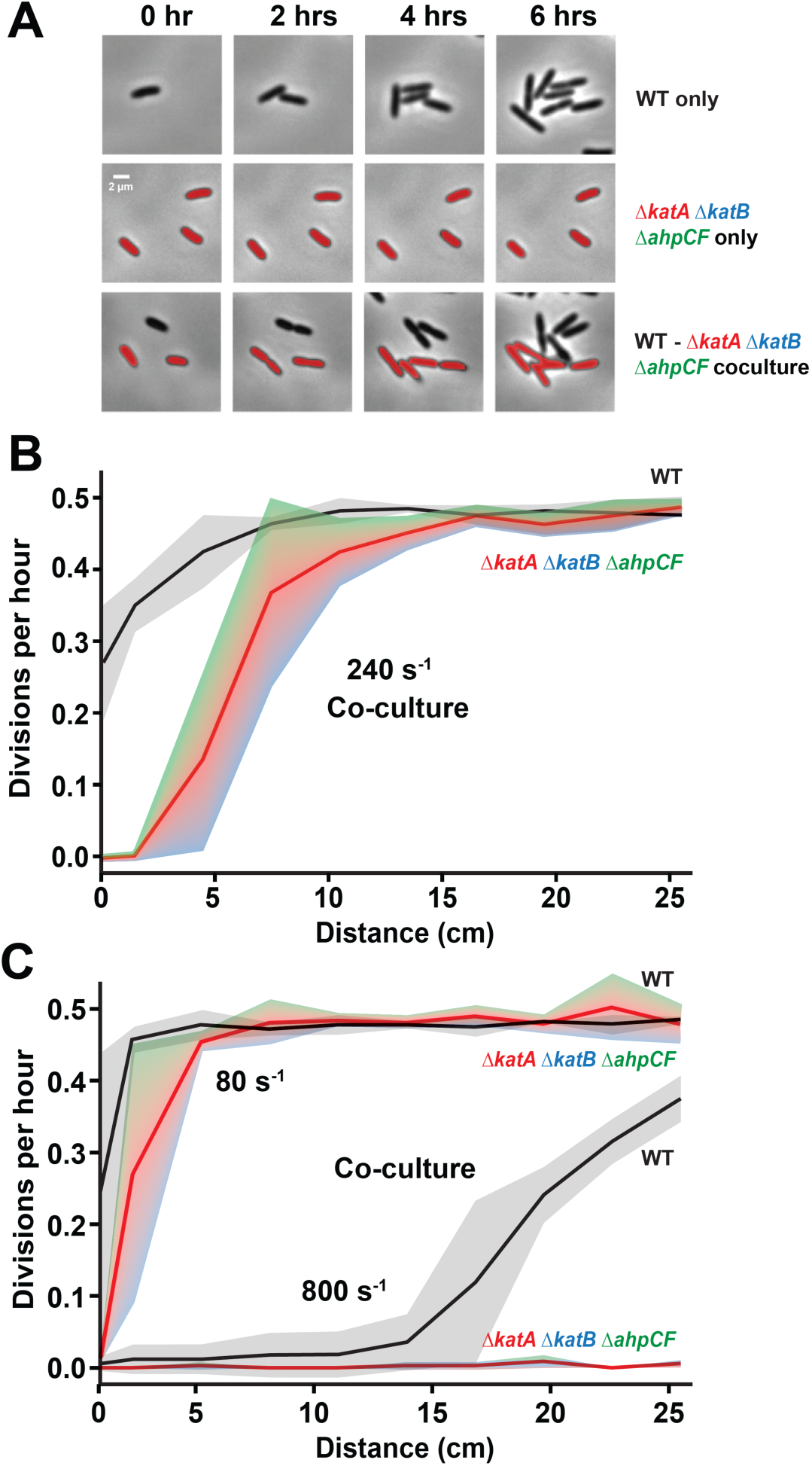
H_2_O_2_ resistant cells protect H_2_O_2_ sensitive cells in co-culture. **(A)** Cell images representative of growth at 13.5 cm into the 27cm channel over 6 hours. Scale bar is 2 µm. **(B)** Quantification of growth at 8 µM H_2_O_2_ in co-culture (90% wild-type and 10% *ΔkatA ΔkatB ΔahpCF* cells). Growth measured at a shear rate of 240 s^-1^. **(C)** Quantification of growth at 8 µM H_2_O_2_ in co-culture (90% wild-type and 10% *ΔkatA ΔkatB ΔahpCF* cells). Growth measured at shear rates of 80 or 800 s^-1^. Quantification shows the average and shaded regions show SD of three biological replicates.

How does flow influence population dynamics in co-culture? As flow can alter the balance between H_2_O_2_ delivery and detoxification, we reasoned that increasing delivery with flow could inhibit population growth. To examine the impact of flow, we measured growth of co-cultured resistant (wild-type) and sensitive cells (*ΔkatA ΔkatB ΔahpCF*) with 8 µM H_2_O_2_ at varied shear rates. At low flow (80 s^-1^), both resistant and sensitive cells grew throughout most of the channel (Figure 5C). At medium flow (240 s^-1^), resistant cells grew throughout most of the channel, while sensitive cells could only grow in the middle and end of the channel (Figure 5B). At high flow (800 s^-1^), resistant cells only grew at the end of the channel, and sensitive cells did not grow at all (Figure 5C). Thus, increasing flow can influence population dynamics by inhibiting the ability of resistant cells to protect sensitive cells. Together, our results demonstrate how physical (flow), chemical (host-relevant H_2_O_2_), and biological (H_2_O_2_ detoxification) factors interact to shape bacterial populations.

## Discussion

Combining host-relevant H_2_O_2_ and shear flow, our results demonstrate how local H_2_O_2_ detoxification provides global protection across bacterial populations. First, we demonstrated that micromolar levels of H_2_O_2_ activate three OxyR-regulated scavenging systems (KatA, KatB, and AhpCF) in *P. aeruginosa* (Figure 1). Second, we used long-channel microfluidic devices to establish a link between H_2_O_2_ scavenging and spatial gradients of growth across *P. aeruginosa* populations (Figure 2). Third, we learned that M9 minimal medium has trace amounts of H_2_O_2_ (∼0.5 µM), which are sufficient to create spatial growth gradients of H_2_O_2_-sensitive *ΔoxyR* mutant cells (Figure 3). Fourth, we used a combination of genetics and biochemistry to establish that KatA, KatB, and AhpCF are each sufficient to detoxify H_2_O_2_ (Figure 4). Fifth, we used co-culture experiments to demonstrate that H_2_O_2_ resistant cells can cross-protect H_2_O_2_ sensitive cells, and that cross protection is inhibited in higher flow regimes (Figure 5). Together, our results highlight how using microfluidics to study bacterial populations under host-relevant conditions can reveal unexpected interactions between physical, chemical, and biological factors.

Individual cells carry out biochemical processes that have population-level consequences. Our work demonstrated that H_2_O_2_ detoxification by individual cells leads to population-level protection (Figure 2), including protection of cells unable to detoxify H_2_O_2_ (Figure 5). We wondered if our results could shed light on the evolutionary pressures driving selection of H_2_O_2_ detoxification. We imagined two hypotheses: H_2_O_2_ detoxification evolved to protect individual cells or H_2_O_2_ detoxification evolved to protect populations. We prefer the hypothesis that H_2_O_2_ detoxification evolved to protect individual cells, and population-level protection occurs as a byproduct. We prefer the individual protection hypothesis because cells have H_2_O_2_ detoxification capacity that is better than necessary for population protection. Specifically, *P. aeruginosa* and other bacterial species (16, 22, 33–35) have evolved multiple scavenging systems that are each sufficient to clear H_2_O_2_ from the environment, indicating that cell entry is the rate limiting step for population-level protection. As individual cells protect themselves better than necessary for population-level protection, we conclude that H_2_O_2_ detoxification is fundamentally a self-defense mechanism that inadvertently performs a de facto public service.

Most laboratory experiments studying the effect of H_2_O_2_ on bacteria use millimolar doses, but host environments typically contain low micromolar concentrations. As bacteria rapidly detoxify H_2_O_2_, it is technically challenging to stress bacteria with host-relevant H_2_O_2_ doses. Here, we used microfluidics to deliver H_2_O_2_ faster than cells could detoxify it, thereby holding H_2_O_2_ levels constant. In a previous microfluidic study, we demonstrated that chemically complex LB medium contained ∼10 µM H_2_O_2_ (13), which triggered H_2_O_2_ stress responses in a flow-dependent manner. Similarly, we showed here that some *P. aeruginosa* mutants struggle to grow on LB plates (Figure S5), reinforcing the conclusion that LB has H_2_O_2_. Based on those observations, we chose to use defined M9 minimal medium for this study, which we assumed did not contain H_2_O_2_. To our surprise, we discovered that M9 minimal medium inhibited growth of our *ΔoxyR* mutant, and that inhibition was abolished by catalase treatment. We subsequently established that our M9 minimal medium contains ∼0.5 µM H_2_O_2_, which likely accumulates due to glucose oxidation (36). Our serendipitous observations establish that low micromolar H_2_O_2_ concentrations stress bacteria and should encourage microbiologists to take a closer look at their media.

“Where in the world do bacteria experience oxidative stress?” (18) Many bacterial pathogens require H_2_O_2_ stress responses for optimal fitness in animal infection models (37–41), indicating that bacteria experience oxidative stress during infection. Host environments typically contain micromolar H_2_O_2_ levels (28–30), which can only transiently stress bacterial populations and don’t appear to inhibit growth unless constantly replenished (11, 36). Thus, we predict that bacteria are primarily inhibited by host-relevant H_2_O_2_ when flow is also present. Additionally, our results help support the conclusion that the balance of flow-based H_2_O_2_ delivery and population-level H_2_O_2_ detoxification is a key determinant of bacterial survival during infection. More broadly, our findings indicate how physical stressors can amplify the biological impact of chemical stressors, highlighting the need to incorporate both host-relevant physical and chemical factors in microbiology experiments.

## Acknowledgements

We thank Gilbert Padron, Jessica Palalay, Piyush Sharma, Evan Johnson, Iota Chen, and Jim Imlay for helpful discussions and comments on the manuscript.

## Contributions

A.S., A.M.S., and J.E.S. designed research. A.S. and A.M.S performed research. A.S., A.M.S., and J.E.S. analyzed data. A.S. and J.E.S. wrote the paper.

## Supplementary Information

### Materials and Methods

#### Strains and growth conditions

The bacterial strains and plasmids used in this paper are described in Supplementary Tables S1 and S2. *P. aeruginosa* cultures were grown in either LB broth or M9 minimal medium on a cell culture roller drum, and on LB plates (1.5% Bacto Agar) at 37°C. LB broth was prepared using LB broth Miller (BD Biosciences). M9 minimal medium was prepared as 1X M9 salts, 0.4% glucose, 2 mM MgSO_4_, and 100 µM CaCl_2_. M9 minimal media supplied with casamino acids was prepared using M9CA Medium Broth, powder (VWR Life Sciences) supplemented with 1 M magnesium sulfate (Sigma), 1 M calcium chloride (Sigma) and 20% glucose (VWR).

#### Generation of *P. aeruginosa* mutants

Gene deletions were generated using the lambda Red recombinase system as previously described (13). Briefly, a deletion construct was assembled using Gibson assembly with three PCR products. First, a segment of approximately 500 bp upstream of the target insertion site was amplified from PA14 genomic DNA. Second, a fragment containing the *aacC1* ORF flanked by FRT sites was amplified from plasmid pAS03. Third, a segment of approximately 500 bp downstream of the target insertion site was amplified from PA14 genomic DNA. The three fragment Gibson product was transformed into PA14 cells expressing the plasmid pUCP18-RedS. Colonies were selected on 30 µg/mL gentamycin, followed by counter-selection of mutants of interest on 5% sucrose. Subsequently, pFLP2 was used to flip out the gentamicin antibiotic resistance gene and the colonies were selected using 300 µg/mL carbenicillin. This resulted in deletion strains with FRT sites.

#### Quantification of RNA levels with qRT-PCR

RNA was isolated from mid-log cells 5 minutes after H_2_O_2_ treatment from 3 biological replicates using TRIzol^®^ Max^TM^ Bacterial RNA Isolation Kit (ThermoFisher). Preheated 200 μL Max Bacterial Enhancement Reagent was used to resuspend the cell pellet in a 1.5 mL Eppendorf tube. After incubation at 95°C for 4 minutes, 1 mL TRIzol was added followed by incubation at room temperature for 5 minutes. Phase separation was performed using 200 µL chloroform followed by centrifugation for 15 minutes at 12,000g, at 4°C. The aqueous phase ∼400 µL was then transferred to a fresh RNAse-free Eppendorf tube. The RNA was then precipitated using cold isopropanol and 75% ethanol. The RNA pellet was then air-dried before resuspending in 50 µL RNase-free water.

Quantitative real-time PCR experiments were performed using Luna® Universal One-Step RT-qPCR Kit (NEB). After thawing the components including Luna Universal One-Step Reaction Mix and primers on ice, the reaction mixture was combined by pipetting. The reaction mix was aliquoted into qPCR wells in the plate (MicroAmp™ Fast Optical 96-Well Reaction Plate, 0.1 mL, ThermoFisher). After adding the RNA template in the respective wells, the plate was sealed using MicroAmp™ Optical Adhesive Film (ThermoFisher). The plate was spun at 3,000 rpm for 1 minute to remove bubbles. The reactions were run on Applied biosystems quantitative PCR machine, and all the acquired Ct values were analyzed using 5S rRNA as an internal control. Primers listed in Table S2 were used.

#### Fabrication of Microfluidic devices

Microfluidic devices were fabricated as previously described (13). Briefly, microfluidic devices were made of Polydimethylsiloxane (Dow SYLGARD 184 Kit) at a 1:10 ratio and plasma-treated to bond on a 60 mm x 35 mm x 0.16 mm superslip micro cover glass (Ted Pella, Inc.). The devices used had channels that were 500 µm wide x 50 µm tall x 27 cm long.

#### Shear rate calculations

Shear rate experienced by cells in microfluidic devices was calculated using this equation:

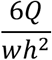

Where *Q* is flow rate, *w* is channel width, and *h* is channel height. The unit for shear rate specified as the inverse of seconds (s^-1^).

#### Phase contrast and fluorescent microscopy

Timelapse images were captured on a Nikon ECLIPSE Ti2-E inverted microscope using the NIS Elements interface. The microscope is equipped with a Nikon 40x Plan Ph2 0.95 NA objective, Hamamatsu ORCA-Flash4.0 LT3 Digital CMOS camera, and Lumencor SOLA Light Engine LED light source.

#### Preparation of microfluidic devices with *P. aeruginosa*

Microfluidic devices were loaded with cells as previously described (13). Briefly, all experiments were performed at approximately 22°C and with mid-log bacterial cultures. Cells were loaded into the microfluidic device and were allowed to settle in the device for 10 minutes prior to exposure to flow. The device set-up involves the use of plastic 5 mL syringes (BD) with attached tubing connecting the needle to the inlet of the device (Brain Tree Scientific Polyethylene Tubing; ID 0.015” x OD 0.043”) that is sheathed over a 26-gauge x 1/2” hypodermic needle (Air-tite Products). These syringes were situated on a syringe pump (KD Scientific Legato 210) which was used to produce fluid flow. The outlet of the device employed the same tubing and vacated into a bleach-containing waste container. The syringe pump was used to generate flow rates of 1-10 µL/min, which correspond to shear rates of 80-800 s^-1^.

#### Quantification of growth of *P. aeruginosa* in microfluidic devices

Single-cell growth was measured in M9 medium with glucose in microfluidic devices. After seeding the devices with mid-log cells, constant M9 minimal medium with defined H_2_O_2_ concentrations was supplied to the microfluidic devices using a syringe pump. Images were taken every 5 minutes up to 6 hours. ImageJ software was used to visualize and track growth, and the number of divisions were measured as the number of divisions per hour in the device. For each biological replicate, 20 cells per field at random were tracked for growth.

#### H_2_O_2_ scavenging assay

H_2_O_2_ scavenging assay was performed using Fluorimetric Hydrogen Peroxide Assay Kit MAK165 (Sigma). *P. aeruginosa* cultures were grown in LB to reach 0.2 OD. Cells were then diluted 1:10 in 1X PBS (phosphate-buffered saline) and were washed twice using 1X PBS. Simultaneously, a master mix of assay buffer, horse radish peroxidase, and red peroxidase substrate was prepared according to the kit protocol. A standard curve was generated using known H_2_O_2_ concentrations. The PBS washed cells were exposed to 10 µM H_2_O_2_, and H_2_O_2_ concentrations were measured overtime for 30 minutes. Equal volumes of cell suspension and master mix (50 µL each) were added to wells of a 96-well plate for each time point.

Fluorescence was measured using a microplate reader at excitation wavelength of 540 nm and emission wavelength of 590 nm. H_2_O_2_ concentrations were calculated using the standard curve at each time point to assess scavenging activity.

---

**Figure S1:**
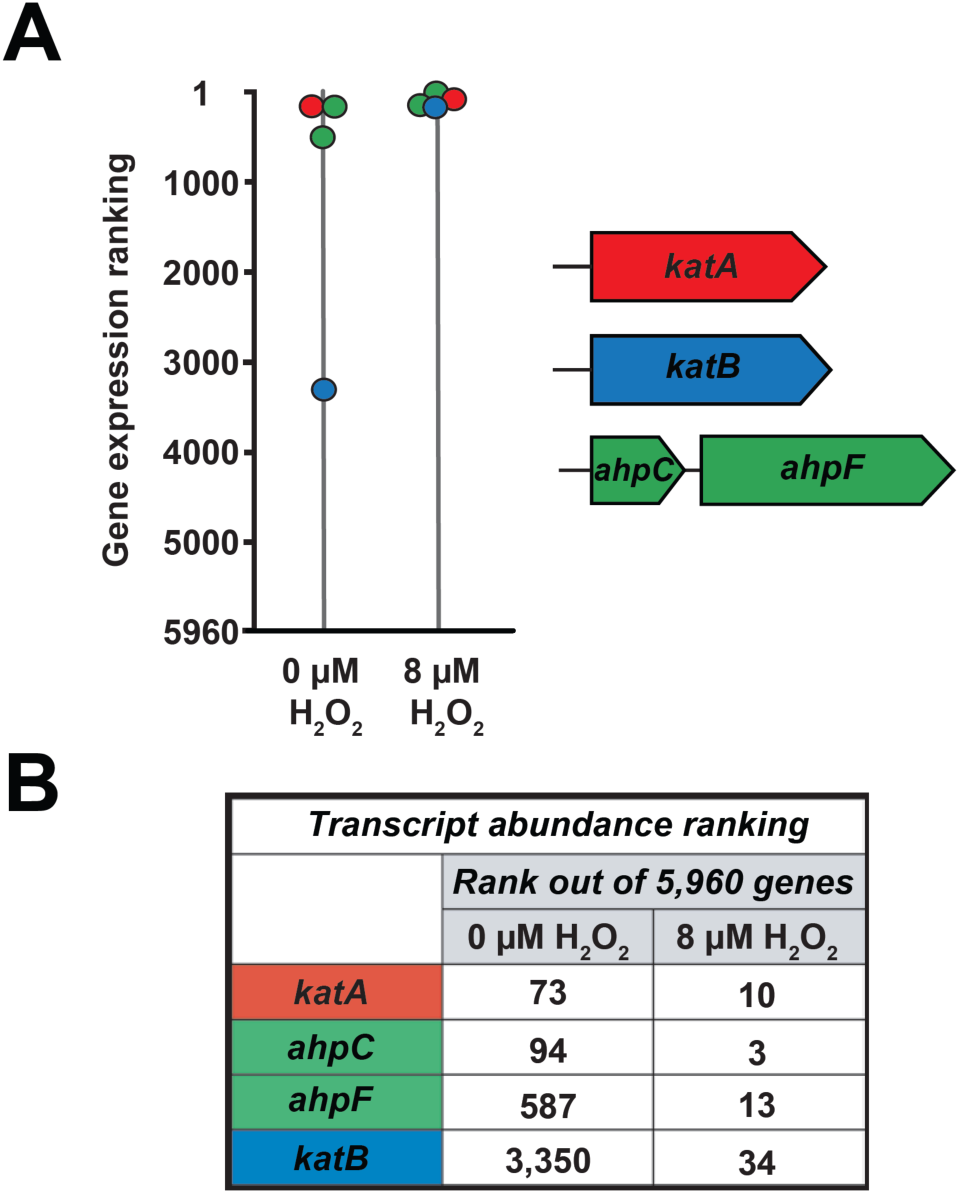
Transcript ranking levels of H_2_O_2_ scavenging genes in response to 8 µM H_2_O_2_. **(A)** Gene expression ranking levels with and without 8 µM H_2_O_2_ exposure for 5 minutes. Higher ranking represents higher RNA levels. **(B)** Transcript abundance levels for H_2_O_2_ scavenging genes *katA, katB, ahpC, ahpF.* (Adapted from Figure 1 of (11))

**Figure S2:**
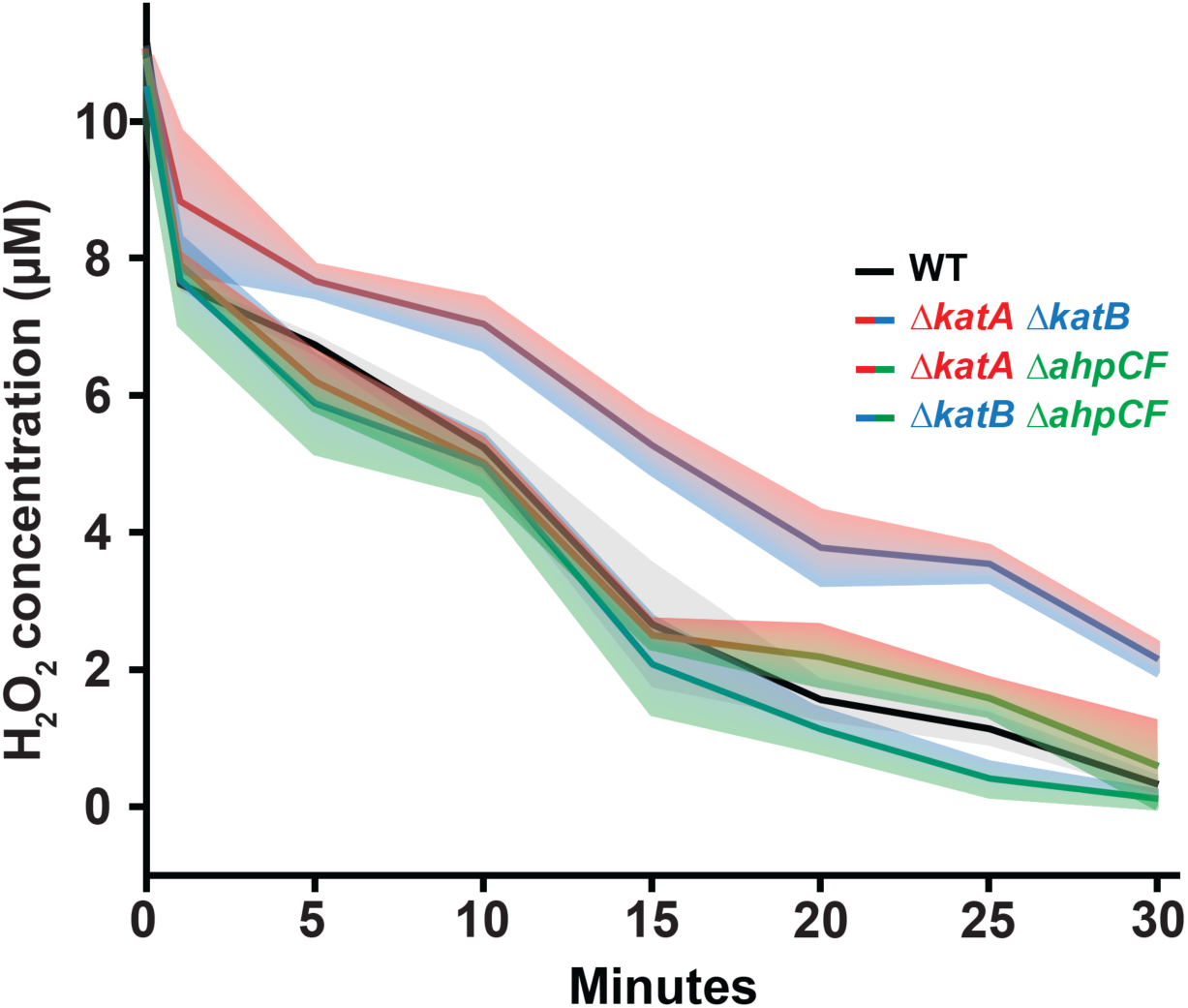
H_2_O_2_ scavenging systems can break down extracellular H_2_O_2_. H_2_O_2_ scavenging by H_2_O_2_ sensitive mutants *ΔkatA ΔkatB*, *ΔkatA ΔahpCF*, *ΔkatB ΔahpCF*. All scavenging assays are performed at a 1:10 dilution of 0.2 OD cells for 30 minutes. Quantification shows the average and shaded regions show SD of three biological replicates.

**Figure S3:**
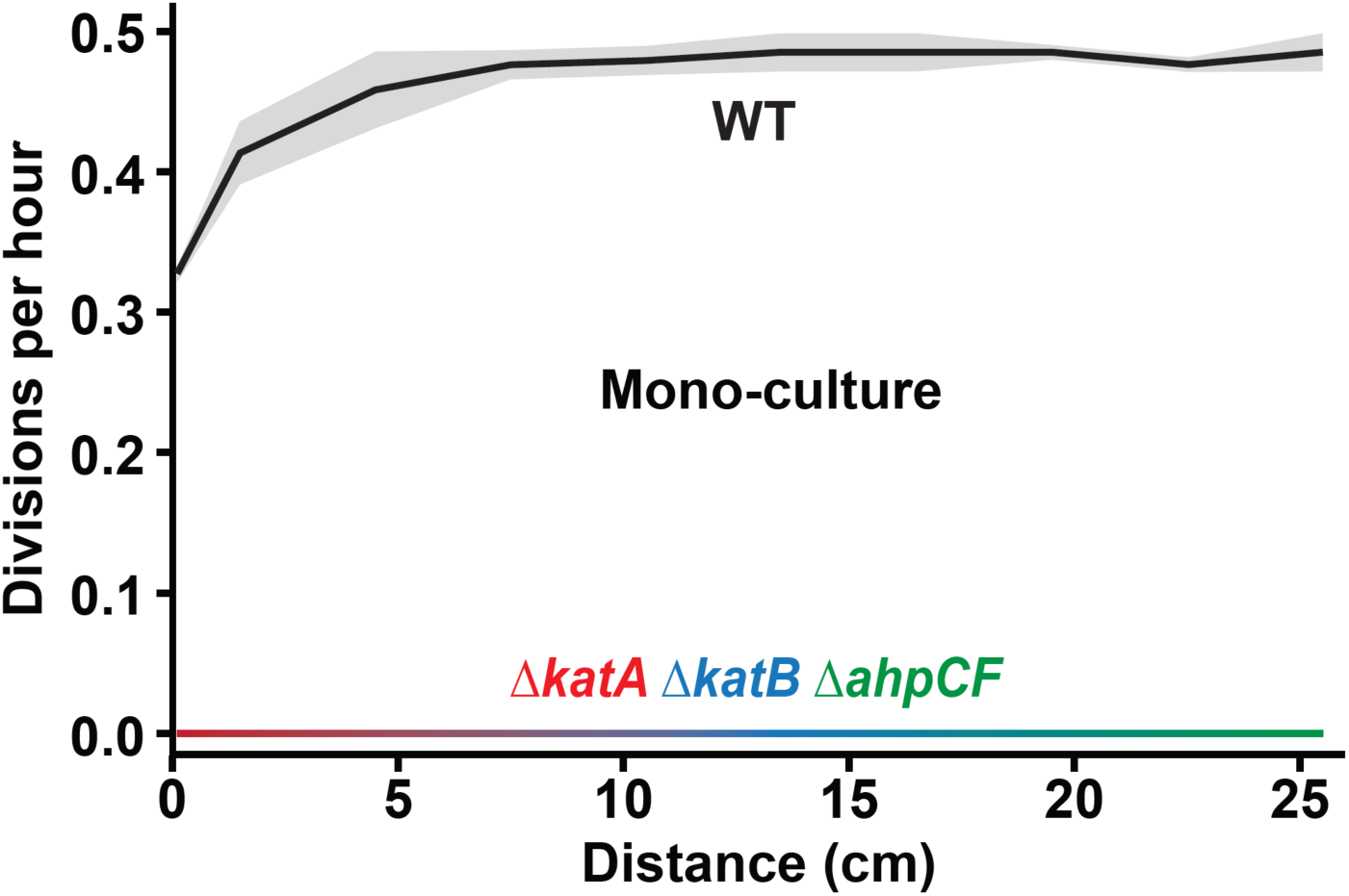
Cells lacking all three scavenging systems do not grow in mono-culture. Quantification of growth at 8 µM H_2_O_2_ for WT and *ΔkatA ΔkatB ΔahpCF* cells in mono-culture. Growth measured at a shear rate of 240 s^-1^. Quantification shows the average and shaded regions show SD of three biological replicates.

**Figure S4:**
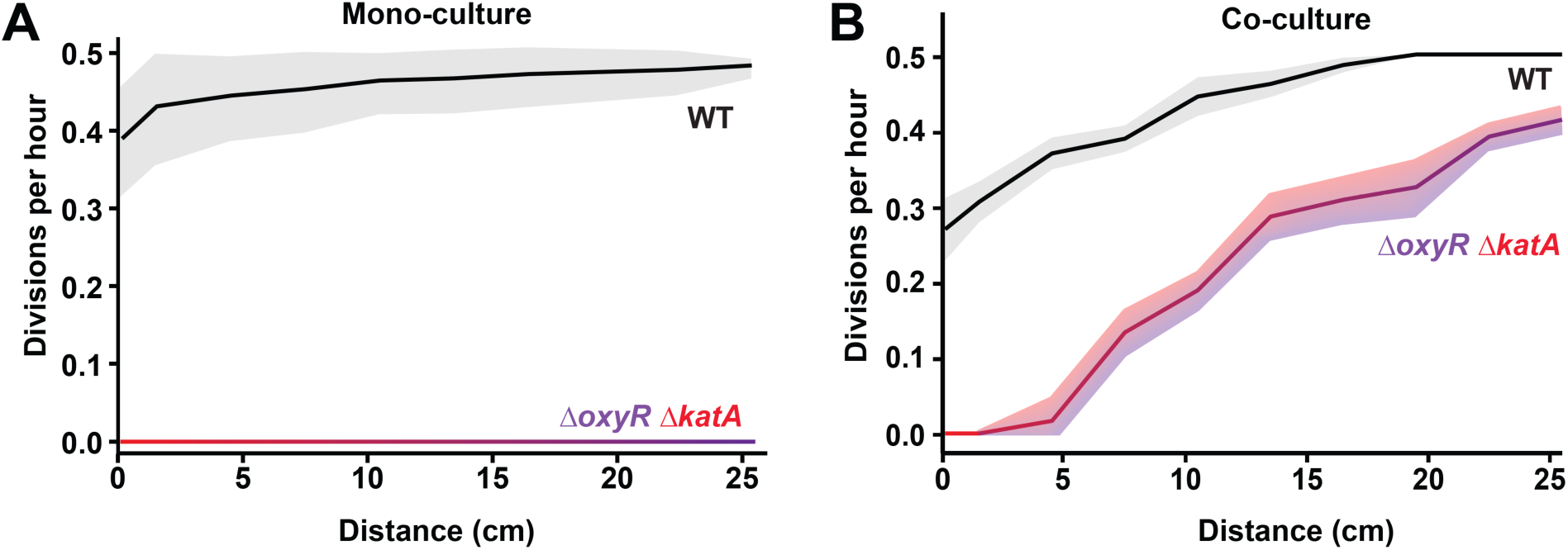
Wild-type cells protect *ΔoxyR ΔkatA* cells in co-culture. **(A)** Quantification of growth at 8 µM H_2_O_2_ in mono-culture. **(B)** Quantification of growth at 8 µM H_2_O_2_ in co-culture (90% wild-type and 10% *ΔoxyR ΔkatA* cells). Growth measured at a shear rate of 240 s^-1^. Quantification shows the average and shaded regions show SD of three biological replicates.

**Figure S5:**
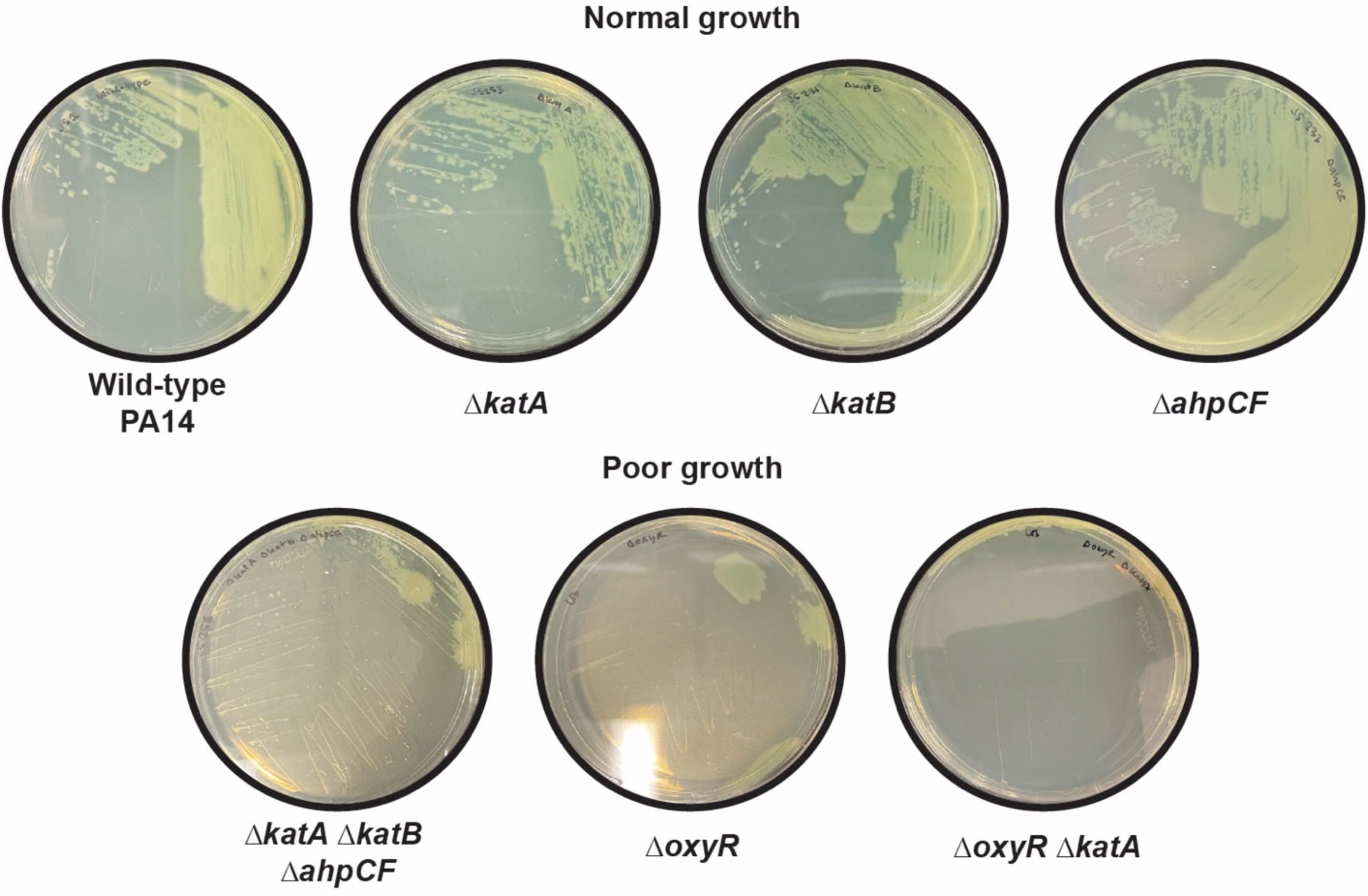
Cells lacking all three scavenging systems or OxyR exhibit poor growth on LB plates. Overnight growth on LB plates for the following *P. aeruginosa* strains: Wild-type (PA14), *ΔkatA*, *ΔkatB*, *ΔahpCF*, *ΔkatA ΔkatB ΔahpCF*, *ΔoxyR*, *ΔoxyR ΔkatA*. As LB generates H_2_O_2_ (13, 36), H_2_O_2_ sensitive mutants (*ΔkatA ΔkatB ΔahpCF*, *ΔoxyR*, *ΔoxyR ΔkatA*) are impaired at growth. Deletion of single scavenging system (*ΔkatA*, *ΔkatB*, or *ΔahpCF*) does not impact growth.

**Table S1:**
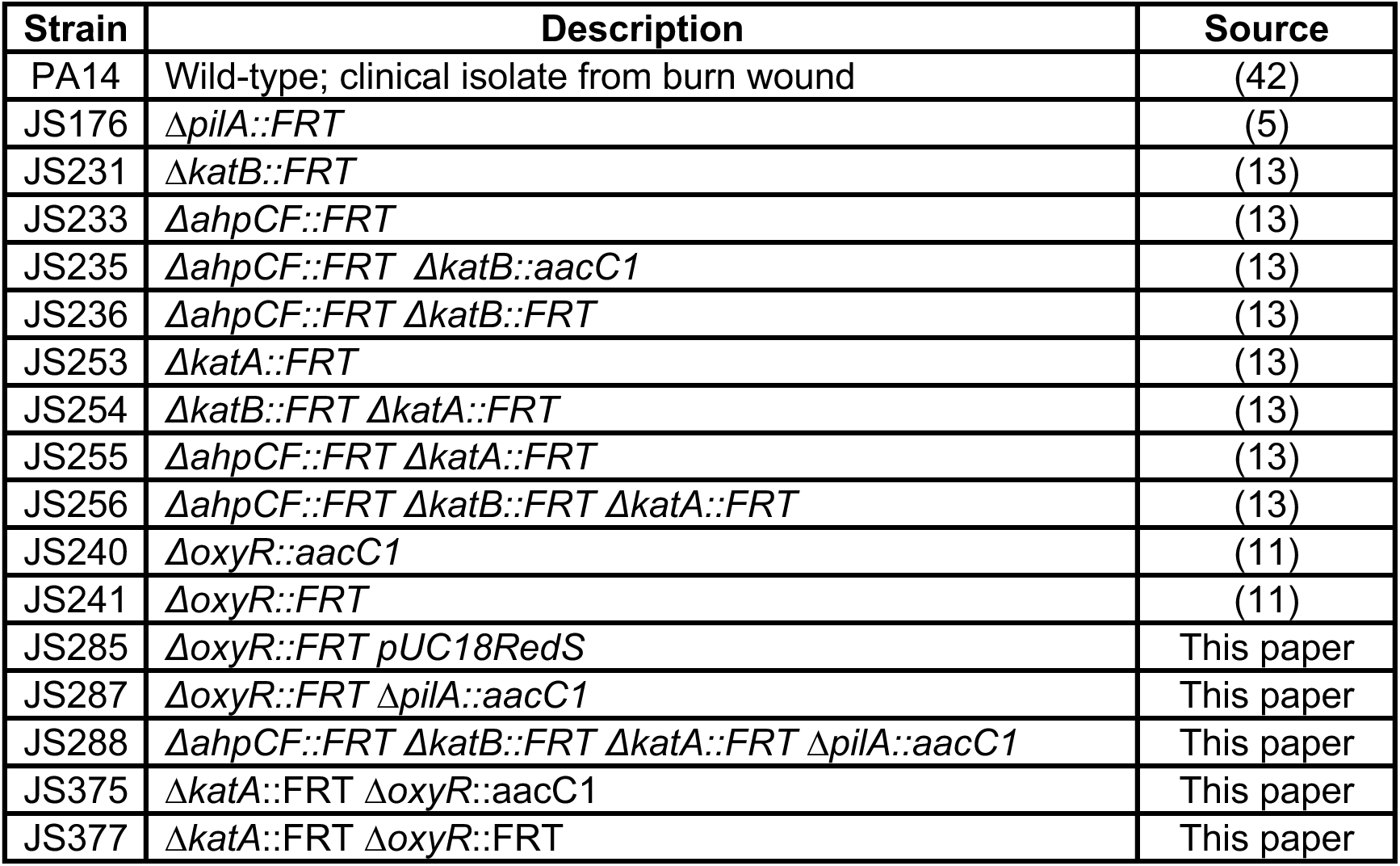
Strains used in the study.

| Table S1: Strains used in the study |  |  |
| --- | --- | --- |
| Strain | Description | Source |
| PA14 | Wild-type; clinical isolate from burn wound | (42) |
| JS176 | $\Delta pilA::FRT$ | (5) |
| JS231 | $\Delta katB::FRT$ | (13) |
| JS233 | $\Delta ahpCF::FRT$ | (13) |
| JS235 | $\Delta ahpCF::FRT \Delta katB::aacC1$ | (13) |
| JS236 | $\Delta ahpCF::FRT \Delta katB::FRT$ | (13) |
| JS253 | $\Delta katA::FRT$ | (13) |
| JS254 | $\Delta katB::FRT \Delta katA::FRT$ | (13) |
| JS255 | $\Delta ahpCF::FRT \Delta katA::FRT$ | (13) |
| JS256 | $\Delta ahpCF::FRT \Delta katB::FRT \Delta katA::FRT$ | (13) |
| JS240 | $\Delta oxyR::aacC1$ | (11) |
| JS241 | $\Delta oxyR::FRT$ | (11) |
| JS285 | $\Delta oxyR::FRT pUC18RedS$ | This paper |
| JS287 | $\Delta oxyR::FRT \Delta pilA::aacC1$ | This paper |
| JS288 | $\Delta ahpCF::FRT \Delta katB::FRT \Delta katA::FRT \Delta pilA::aacC1$ | This paper |
| JS375 | $\Delta katA::FRT \Delta oxyR::aacC1$ | This paper |
| JS377 | $\Delta katA::FRT \Delta oxyR::FRT$ | This paper |

**Table S2:**
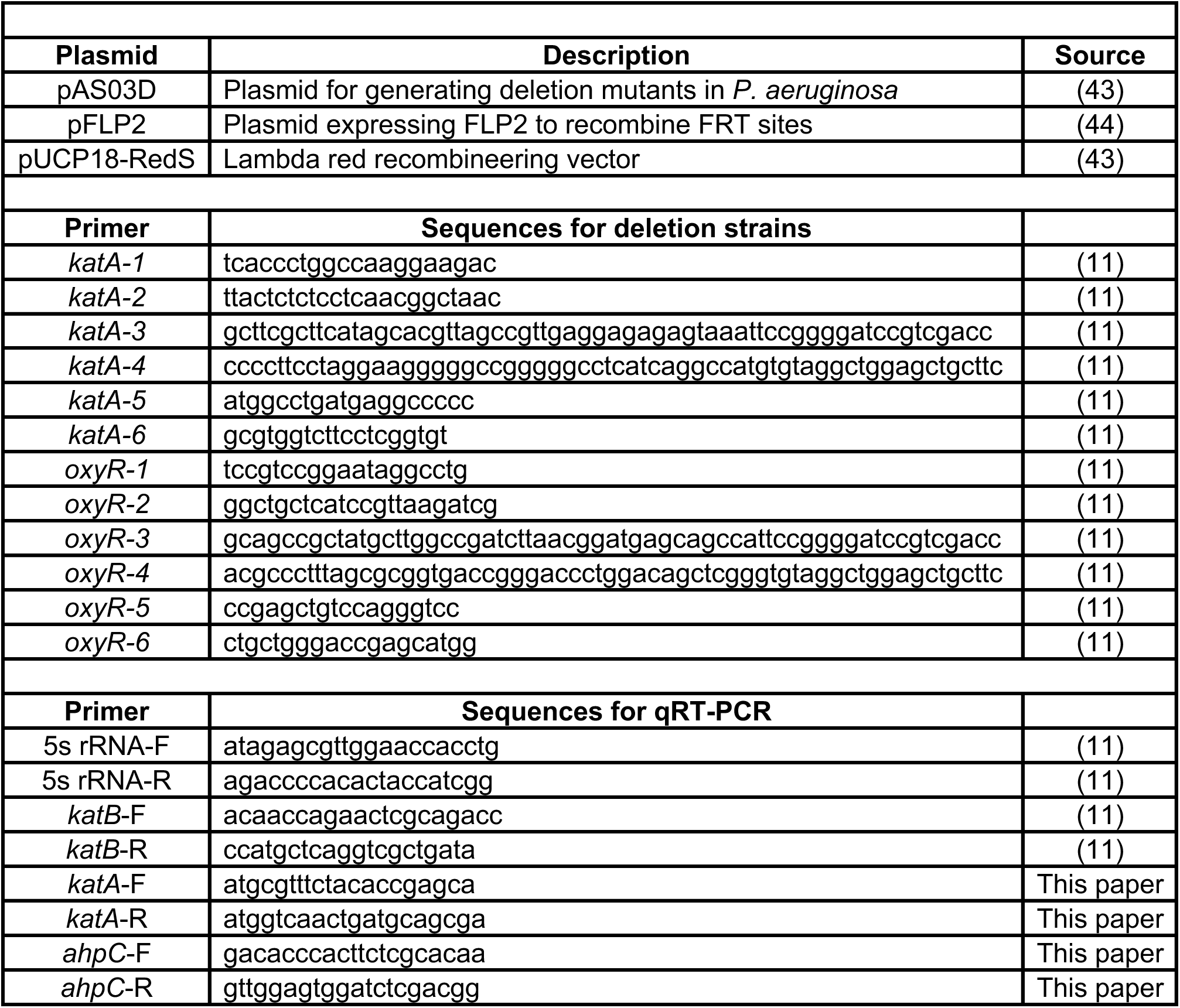
Plasmids and primers used in the study.

